# The FDA-approved drug Alectinib compromises SARS-CoV-2 nucleocapsid phosphorylation and inhibits viral infection in vitro

**DOI:** 10.1101/2020.08.14.251207

**Authors:** Tomer M. Yaron, Brook E. Heaton, Tyler M. Levy, Jared L. Johnson, Tristan X. Jordan, Benjamin M. Cohen, Alexander Kerelsky, Ting-Yu Lin, Katarina M. Liberatore, Danielle K. Bulaon, Edward R. Kastenhuber, Marisa N. Mercadante, Kripa Shobana-Ganesh, Long He, Robert E. Schwartz, Shuibing Chen, Harel Weinstein, Olivier Elemento, Elena Piskounova, Benjamin E. Nilsson-Payant, Gina Lee, Joseph D. Trimarco, Kaitlyn N. Burke, Cait E. Hamele, Ryan R. Chaparian, Alfred T. Harding, Aleksandra Tata, Xinyu Zhu, Purushothama Rao Tata, Clare M. Smith, Anthony P. Possemato, Sasha L. Tkachev, Peter V. Hornbeck, Sean A. Beausoleil, Shankara K. Anand, François Aguet, Gad Getz, Andrew D. Davidson, Kate Heesom, Maia Kavanagh-Williamson, David Matthews, Benjamin R. tenOever, Lewis C. Cantley, John Blenis, Nicholas S. Heaton

**Affiliations:** Meyer Cancer Center, Weill Cornell Medicine, New York, NY, USA; Department of Medicine, Weill Cornell Medicine, New York, NY, USA; Englander Institute for Precision Medicine, Institute for Computational Biomedicine, Weill Cornell Medicine, New York, NY, USA; Department of Physiology and Biophysics, Weill Cornell Medicine, New York, NY, USA; Tri-Institutional PhD Program in Computational Biology & Medicine, Weill Cornell Medicine/Memorial Sloan Kettering Cancer Center/The Rockefeller University, New York, NY, USA; Department of Molecular Genetics and Microbiology, Duke University School of Medicine, Durham, NC, USA; Cell Signaling Technology, Danvers, MA, USA; Department of Microbiology, Icahn School of Medicine at Mount Sinai, New York, NY, USA; The Biochemistry, Structural, Developmental, Cell and Molecular Biology Allied PhD Program, Weill Cornell Medicine, New York, NY, USA; Division of Gastroenterology and Hepatology, Department of Medicine, Weill Cornell Medicine, 1300 York Ave, New York, NY, USA; Department of Surgery, Weill Cornell Medicine, 1300 York Ave, New York, USA; Department of Dermatology, Weill Cornell Medicine, New York, NY, USA; Department of Microbiology and Molecular Genetics, Chao Family Comprehensive Cancer Center, University of California Irvine School of Medicine, Irvine, CA, USA; Department of Cell Biology, Duke University School of Medicine, Durham, NC, USA; Broad Institute of MIT & Harvard, Cambridge, MA, USA; Department of Pathology, Harvard Medical School, Cambridge, MA, USA; Cancer Center and Department of Pathology, Massachusetts General Hospital, Boston, MA, USA; School of Cellular and Molecular Medicine, University of Bristol, Bristol, BS8 1TD, UK; Proteomics Facility, University of Bristol, Bristol, BS8 1TD, UK; Department of Pharmacology, Weill Cornell Medicine, New York, NY, USA; Department of Biochemistry, Weill Cornell Medicine, New York, NY, USA; Duke Human Vaccine Institute, Duke University School of Medicine Durham, NC, USA; Duke Cancer Institute, Duke University School of Medicine, Durham, NC, USA

## Abstract

While vaccines are vital for preventing COVID-19 infections, it is critical to develop new therapies to treat patients who become infected. Pharmacological targeting of a host factor required for viral replication can suppress viral spread with a low probability of viral mutation leading to resistance. In particular, host kinases are highly druggable targets and a number of conserved coronavirus proteins, notably the nucleoprotein (N), require phosphorylation for full functionality. In order to understand how targeting kinases could be used to compromise viral replication, we used a combination of phosphoproteomics and bioinformatics as well as genetic and pharmacological kinase inhibition to define the enzymes important for SARS-CoV-2 N protein phosphorylation and viral replication. From these data, we propose a model whereby SRPK1/2 initiates phosphorylation of the N protein, which primes for further phosphorylation by GSK-3α/β and CK1 to achieve extensive phosphorylation of the N protein SR-rich domain. Importantly, we were able to leverage our data to identify an FDA-approved kinase inhibitor, Alectinib, that suppresses N phosphorylation by SRPK1/2 and limits SARS-CoV-2 replication. Together, these data suggest that repurposing or developing novel host-kinase directed therapies may be an efficacious strategy to prevent or treat COVID-19 and other coronavirus-mediated diseases.

## INTRODUCTION

In December 2019, a novel human coronavirus, now known as SARS-CoV-2, emerged and began causing a human disease termed COVID-19 (*1, 2*). Since then, a global pandemic has infected countless numbers of people and caused more than a million deaths to date. Due to the prevalence and severity of this disease, the development of therapeutic interventions is of the highest importance. Much attention has been focused on targeting viral proteins and their associated enzymatic activities. In particular, the virally encoded RNA-dependent RNA polymerase (RdRp) and the viral proteases, are attractive potential targets. Remdesivir, the only FDA-approved antiviral for SARS-CoV- 2, is a nucleoside analogue which targets the viral RdRp and causes premature termination of transcription (*3*). Efficacy of this treatment however, unfortunately, appears limited (*4*).

In addition to targeting viral proteins directly, other antiviral development strategies attempt to target host factors that the virus requires to complete its lifecycle. Relative to their hosts, viruses have dramatically less coding space in their genomes and therefore utilize the host to enable means to render virus proteins to be multifunctional. The major advantages of inhibiting a virus indirectly via an essential host factor are two-fold: (1) many viruses may utilize the same host protein, therefore host-directed therapeutics have the potential to be broadly acting and (2) while direct targeting of the virus can rapidly select for resistant viral mutants, it is thought to be much more difficult for a viral mutation to overcome inhibition of a co-opted host protein. While not all host factors are easily targetable, some enzymes such as protein kinases, for which inhibitors have been developed and tested for activity against other diseases such as cancer, are of high interest for host-directed antivirals.

In this report, we focused on defining the kinases that mediate phosphorylation of the SARS-CoV-2 nucleocapsid (N) protein, specifically its SR-rich domain, due to its high level of conservation across coronaviruses, previous data showing that it is highly phosphorylated (*5–10*), and also the understanding that N protein phosphorylation is important for its functionality (*11–16*). After performing phosphoproteomics analysis on both human and monkey cells to identify the phosphorylation sites on the N protein, we utilized high-throughput kinase substrate specificity mapping and *in vitro* phosphorylation assays to define not only which host kinases can phosphorylate the N protein, but also the order of action and the specific phosphorylation sites of each kinase. We then used both genetic knockdown and pharmacological targeting of the key kinases SRPK1/2 to verify their requirement during the replication of multiple human coronaviruses including SARS-CoV-2. Finally, we showed that several SRPK1/2 inhibitors including FDA-approved kinase inhibitor Alectinib, which can inhibit SRPK1/2 (*17*), can compromise SARS-CoV-2 replication in multiple cell lines and primary human pneumocytes. Thus, not only are the enzymatic activities of SRPK1/2 essential for the replication of multiple human coronaviruses via phosphorylation of the N protein, but targeting these kinases may represent an immediate and effective therapeutic strategy to combat coronavirus mediated diseases, including COVID-19.

## RESULTS

In order to first identify the phosphorylation sites on the SARS-CoV-2 nucleocapsid (N) protein with high confidence, we infected both human A549 lung epithelial cells expressing the ACE2 receptor (A549-ACE2) and African green monkey [*Chlorocebus sabaeus*] kidney cells (Vero E6) with SARS-CoV-2, as these lines are highly susceptible to infection by SARS-CoV-2 (*18*). Infected and control cells were harvested in biological triplicate followed by global proteomics and phosphoproteomics analysis using LC-MS (**Figure 1A**). Analysis of the SARS-CoV-2 N protein revealed 14 phosphorylation sites in A549-ACE2 cells, 11 of them specifically in the SR-rich domain (**Table S1**). In Vero cells, 26 phosphorylation sites were detected on the N protein, 15 of them in the SR-rich domain (**Table S1**). Most of the sites in the SR-rich domain were found in both cell lines in our study, as well as in 3 previous phosphoproteomics studies (*8–10*) (**Figure 1B**). To investigate the evolutionary conservation of the SARS-CoV-2 N protein, we next compared the nucleocapsid proteins from 82 different coronaviruses (**Table S2**) and determined the percentage conservation of each amino acid. Interestingly, we noticed that the SR-rich domain is significantly more conserved than the linker domain, and as conserved as the two known functional domains of the N protein – the N-Terminus Domain (NTD, involved in RNA binding (*19–22*)) and the C-Terminus Domain (CTD, involved in protein oligomerization and RNA binding (*23–25*) (**Figure 1B**).

**Figure 1.**
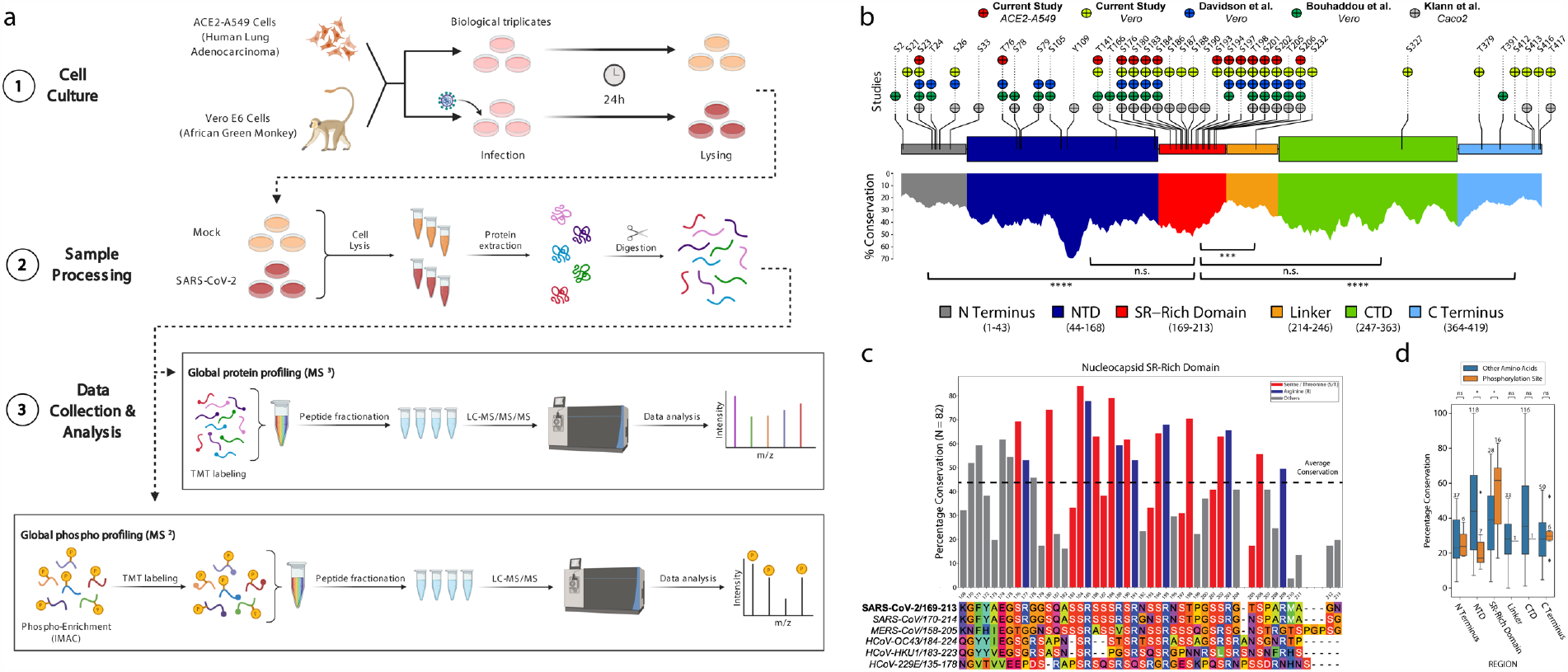
SARS-CoV-2 nucleocapsid protein is heavily phosphorylated at the SR-rich domain. A. Diagram of the proteomics and phosphoproteomics workflow for cells infected with SARS-CoV-2. A549-ACE2 and Vero E6 cells were infected with SARS-CoV-2 (MOI 0.5) and mock infected for 24h (in biological triplicates each). Then, cells were harvested and lysed, and proteins were cleaved into peptides using trypsin. Global protein profiling was carried out using liquid chromatography mass spectrometry (LC-MS) on an aliquot of the peptides from each sample. The rest of the peptides were enriched for phosphorylation using IMAC, and were analyzed by LC-MS as well. Peptides and phosphorylation sites were mapped to the human proteome and the SARS-CoV-2 proteins. B. Phosphorylation sites on the SARS-CoV-2 N protein identified in 5 different phosphoproteomics analyses (two in the current study and three in previously published studies). Most of the sites at the SR-rich domain were detected in at least two independent studies (top). Evolutionary conservation analysis of the different domains of the N protein across 82 different coronaviruses shows that the SR-rich domain is more conserved than the linker domain, and is as conserved as the other two functional domains of the N protein – NTD and CTD (bottom). Mann-Whitney U test: n.s. – not significant, *p<0.05, **p<0.01, ***p<0.001, ****p<0.0001. C. Alignment of the SR-rich domain of the N protein across 6 different human coronaviruses (bottom), and the percentage identity of each amino acid across 82 coronaviruses in multiple species (top). The most conserved residues are serines, threonines and arginines. D. The conservation of the phosphorylation sites in each domain was compared to the conservation of the other amino acids in that domain. Only in the SR-rich domain the phosphorylation sites are significantly more conserved than the rest of the residues. The numbers of amino acids compared in each domain are annotated above the boxes. Mann-Whitney U test: n.s. – not significant, *p<0.05.

Further evolutionary examination of the SR-rich domain shows that the most conserved amino acids in that domain are serines (S), threonines (T) and arginines (R) (**Figure 1C**), indicating that these are the key residues in that domain. Together with previous evidence that the phosphorylation of the SR-rich domain is important for the life cycle of coronaviruses in general (*7, 26*) and SARS-CoV (*5, 11, 13, 15, 27*) in particular, we hypothesized that the phosphorylation of the serines and threonines in this domain is important for viral life cycle of SARS-CoV-2, and that the arginines are likely to be essential for the phosphorylation to occur as well. Finally, comparison of the conservation of the detected phosphorylation sites in each domain to all the other amino acids in that domain shows that only in the SR-rich domain, the phosphorylation sites are significantly more conserved relative to the rest of the region (**Figure 1D**). These data taken together suggest that not only is the N protein highly phosphorylated, but that the phosphorylation of the SR-rich domain in particular may be functionally important for SARS-CoV-2 life cycle.

We next identified the host kinase(s) that phosphorylate the serines and threonines in the SR-domain. The ability of protein kinases to phosphorylate substrates is strongly dependent on the serine/threonine phosphoacceptor’s surrounding amino acid sequence. The majority of human kinases investigated show distinct preferences for or against amino acids surrounding their phosphoacceptor. This is collectively referred to as their *substrate motifs* and is useful for identifying biological substrates. To obtain the substrate motif of a kinase, ours and other laboratories have developed an unbiased approach using combinatorial peptide substrate libraries (*28–31*).

Previous studies have reported that the GSK-3 and SRPK families are involved in the phosphorylation of the SR-rich domain of the nucleocapsid protein of SARS-CoV (*5, 15*). Recent reports have suggested these kinase families to be involved in the phosphorylation of SARS-CoV-2 N protein phosphorylation as well (*32–34*), however an exact phosphorylation model has never been validated experimentally. To that end, we empirically characterized the biochemical substrate specificities of GSK-3α/β and SRPK1/2/3 (**Figure 2A and Figure S1**). Consistent with previous reports (*35–37*), GSK- 3 prefers to phosphorylate serines and threonines that have an already phosphorylated serine or threonine four residues apart toward the C-terminus (designated position +4, relative to the phosphoacceptor). This phenomenon is called *phospho-priming* – in which an already phosphorylated residue promotes phosphorylation of another proximate residue. The SR-rich domain of the N protein contains three chains of serines/threonines regularly repeating every fourth residue: S206-S186, T205-S193 and S188-S176. We therefore that the C-terminal serines/threonines of these chains (the priming sites S206, T205, and S188) are phosphorylated first, which initiates a phosphorylation cascade by GSK-3, resulting in the full phosphorylation of the entire chains.

**Figure 2.**
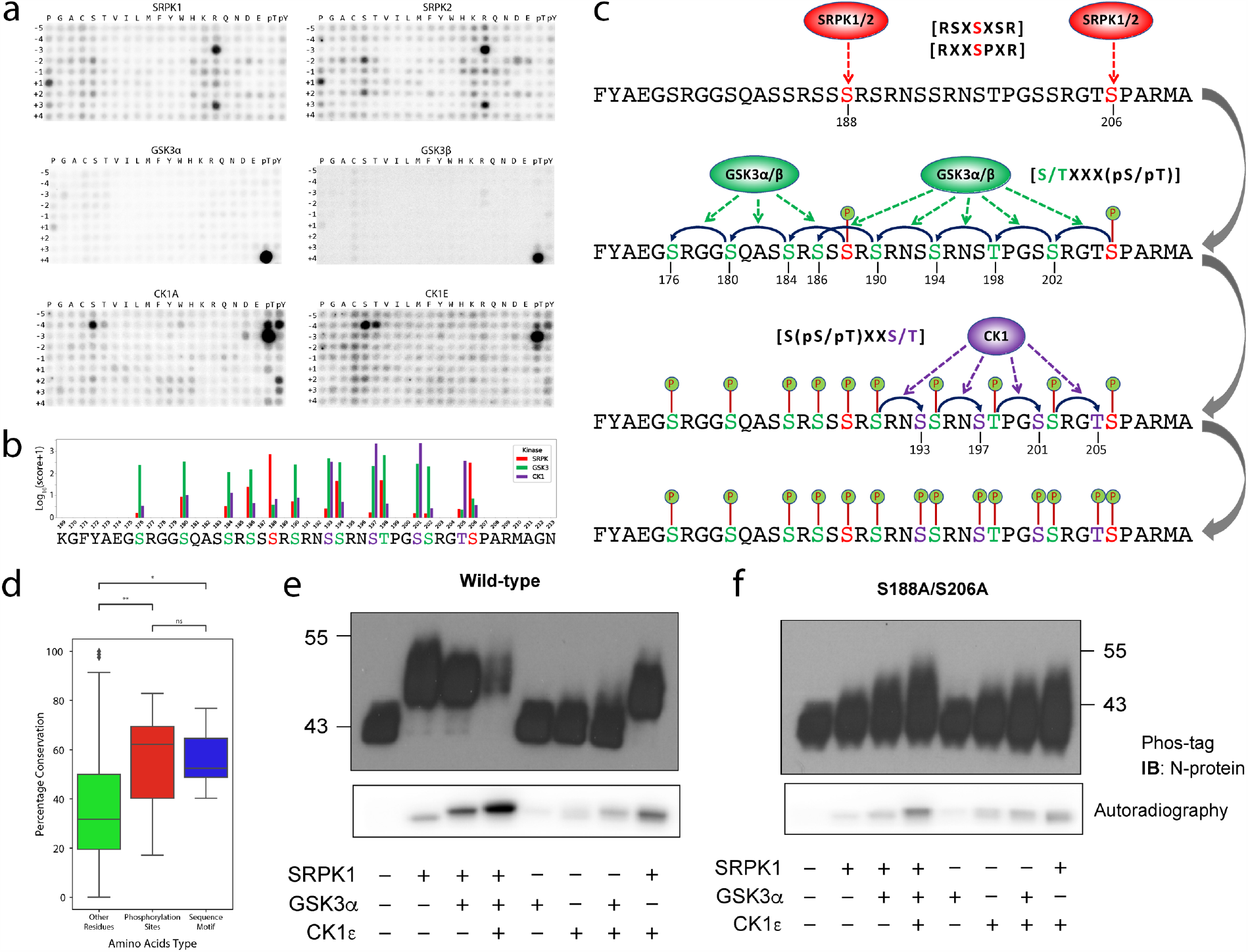
Phosphorylation model of the N protein SR-rich domain by SRPK, GSK-3 and CK1. A. Biochemical substrate specificity of SRPK1/2 (top), GSK-3α/β (middle) and CK1A/G (bottom). SRPKs are selective for arginines at the −3 and +3 positions, serines at the −2 and +2 positions, and proline at the +1 position. GSK-3s are selective for phosphoserine or phosphothreonine at position +4. CK1s are selective for phosphoserine and phosphothreonine at position −3 and for serine at position −4. (see Supplementary Figure 1 for substrate specificities of additional kinases from these families). B. Favorability scores for the different phosphorylation sites at the SR-rich domain according to the SRPK (SRPK1/2/3), GSK-3 (GSK-3α/β) and CK1 (CK1A/D/ε/G1) families. SRPK family scores the highest for S206 and S188, and once phosphorylated, those sites prime GSK-3 family to phosphorylate every serine or threonine located 4 positions toward the N terminus (phosphorylation chains). Finally, CK1 family scores the highest for T205 where both GSK-3 and CK1 families score high for the other sites in its corresponding phosphorylation chain. C. Proposed scheme for the multisite phosphorylation of the nucleocapsid SR-rich domain. Simplified substrate consensus motifs are shown here in parenthesis, whereas detailed logos are provided in Figure S2. D. Evolutionary conservation across 82 different coronaviruses from multiple species shows that the amino acids predicted to be essential for substrate specificity of the priming sites are as conserved as the phosphorylation sites themselves, and are more conserved than the other amino acids in the SR-rich domain. Mann-Whitney U test: n.s. – not significant, *p<0.05, **p<0.01. E. Top: Immunoblot of recombinant N protein on Phos-tag gel after treatment with recombinant kinases SRPK1, GSK-3α, and/or CK1ε. Bottom: SDS-PAGE/autoradiography of recombinant N protein after treatment with SRPK1, GSK-3α, and/or CK1ε, in the presence of ATP[γ-^32^P]. F. Removing S188 and S206 reduced the phosphorylation level of the N protein. Phos-tag gel analysis (top) and autoradiography (bottom) of recombinant N protein with the priming phosphorylation sites mutated (S188A, S206A), performed as described in (E). For comparison, autoradiography of the mutant and WT N proteins was measured together.

In order to match each site to its most likely upstream kinase based on the characterized substrate specificity matrices, we computed a favorability score (see Materials and Methods) for each characterized kinase, for every phosphorylation site in the three phosphorylation chains above (**Figure 2B and Table S2**). Our algorithm predicted S206 and S188 to be phosphorylated by SRPKs (SRPK1/2/3), which have strong preference for arginine at the −3 and +3 positions, serine at the −2 and +2 positions, and proline at the +1 position (**Figure 2A and Figure S1**). Once S206 and S188 are phosphorylated and thus. primed, our algorithm predicted that, as expected, a GSK-3 family member (GSK-3α/β) would sequentially phosphorylate the chain of serine/threonine every 4 residues toward the N-terminus (S202-T198-S194-S190-S186 and S184-S180-S176, respectively).

Finally, since neither the SRPK or GSK-3 families score favorably for the third priming site (T205), we considered additional kinase(s) that might carry this out. Assuming that S202 is phosphorylated by GSK-3 as discussed above, we searched for kinases with strong preference for phosphoserine or phosphothreonine at position −3 as likely kinases to phosphorylate S205. The CK1 family is a second group of phosphopriming-dependent protein kinases that favors strongly phosphoserine or phosphothreonine at position −3 and partially select for unmodified serine at the −4 position (**Figures 2A and Figure S1**). In parallel, our algorithm predicted T205 to be a likely phosphorylation site for CK1, when primed by phosphorylation at S202 (**Figure 2B and Table S2**). Subsequent phosphorylation at S201, S197 and S193 can be carried out either by CK1 or GSK-3. **Figure 2C** summarizes our proposed model for the cluster of phosphorylation sites in the SR-rich domain of the SARS-CoV-2 nucleocapsid.

Interestingly, the amino acids that are dominant in the substrate motif of SRPK (S188: R185/S186/S190/R191; S206: R203/P207/R209) are as conserved as the phosphorylated residues and across coronaviruses are more conserved than the other amino acids in that region, suggesting that those amino acids indeed play an important role in directing the appropriate protein kinases to phosphorylate this region of the N protein (**Figure 2D**).

To test our sequential phosphorylation model, recombinant SARS-CoV-2 N protein was purified and subjected to *in vitro* phosphorylation assays with recombinant SRPK1, GSK- 3α, and CK1ε. Phos-tag gel analysis of the reactions showed an upward shift of the N protein band following treatment with SRPK1, indicating stoichiometric phosphorylation at one or more sites of the N protein. Adding GSK-3 or CK1 without prior treatment with SRPK1 had only modest effects on the phos-tag shift. To determine the amount of phosphate incorporation into N protein, radioactively labeled ATP was included in the phosphorylation reactions and autoradiography of N protein was measured on SDS-PAGE. Treatment with SRPK, GSK-3 and CK1 increased phosphorylation of N protein to a greater extent than the sum of the individual kinase reactions, consistent with our model where SRPK primes the SR-region for phosphorylation by GSK-3 and CK1 (**Figure 2E**). Importantly, adding all three kinases caused a major upward shift in the Phos-tag gel and a reduced detection of protein, suggesting that the highly phosphorylated protein did not efficiently enter the gel (**Figure 2E**). The phospho-null double mutant (S188A, S206A) abolished the phos-tag shift caused by SRPK and showed reduced radioactive incorporation by SRPK, GSK-3 and CK1 (**Figure 2F**). This is consistent with our model that phosphorylation of N protein at S188/S206 by SRPK is the critical priming event for extensive phosphorylation of the SR-rich domain.

We next asked how loss of SPRK proteins would affect viral replication. It has been reported that SRPK1 is expressed at higher amounts in A549 cells relative to SRPK2; we therefore targeted SRPK1 in our ACE2-A549 cells (**Figure S3**) via RNAi (**Figure 3A**). Both the kinase RNA and protein were significantly reduced after treatment (**Figure 3B,C**), and viral RNA levels were correspondingly suppressed (**Figure 3D**). As an orthogonal approach to validate the requirement of SRPK1 during SARS-CoV-2 infection, we utilized the SRPK inhibitor SPHINX31 to block SRPK1/2 activity (*38*). After treatment of A549-ACE2 cells with low micromolar concentrations of SPHINX31, which inhibit both SRPK1 and SRPK2, we observed inhibition of SARS-CoV2 replication (**Figure 3E,F**). We also utilized a second SRPK1/2 inhibitor, SRPIN340 (*39*), and as expected, treatment decreased viral RNA and infectious viral titer in a dose dependent manner at concentrations that were well tolerated by the cells (**Figure 3G,H**).

**Figure 3.**
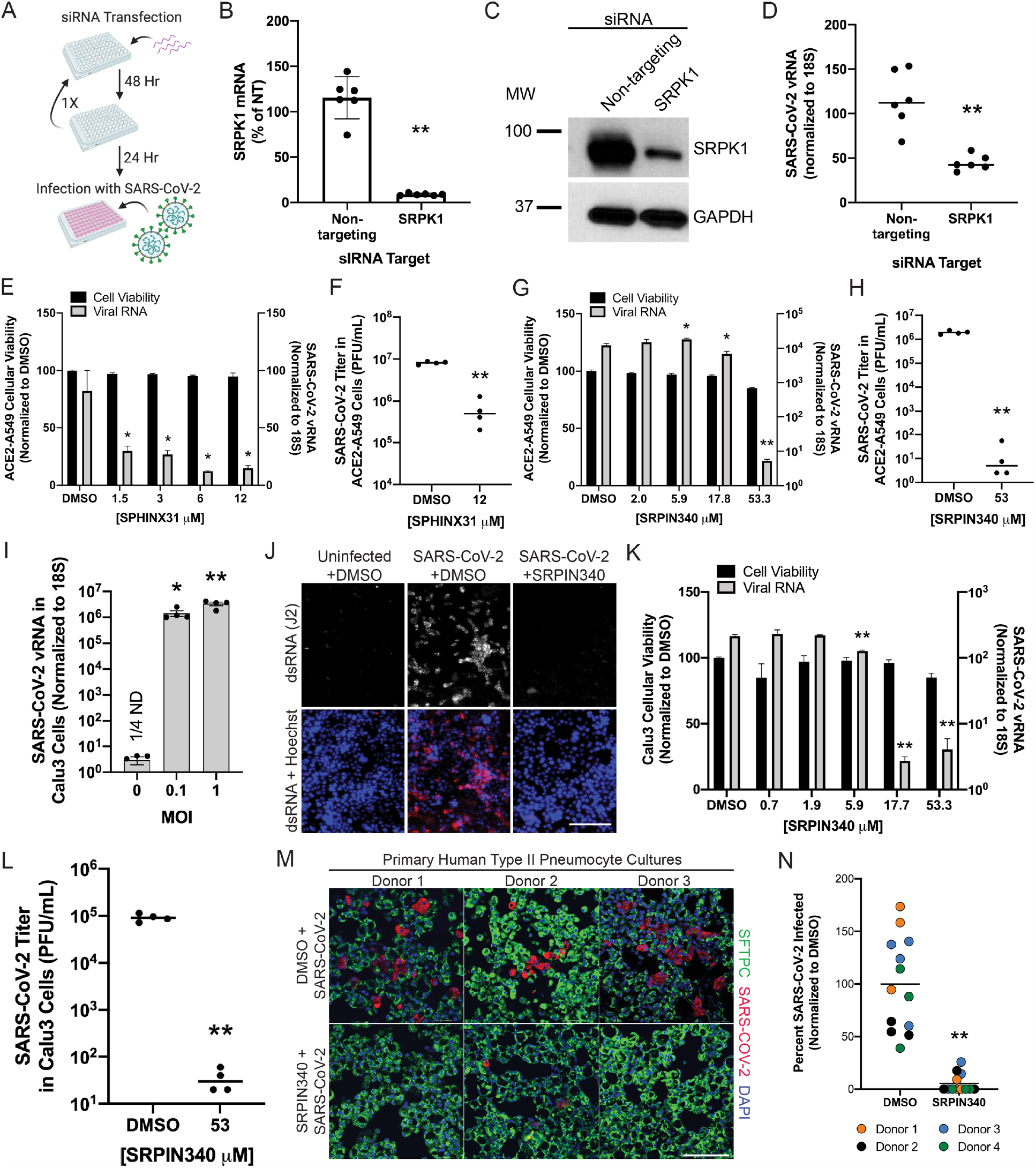
SRPK1/2 inhibitors can suppress SARS-CoV-2 infection. A. Diagram of siRNA transfection. Cells were plated and treated with siRNA on day 0. After 48hrs of transfection, cells were trypsinized again and replated in 24 well plates and transfected with siRNA once again. 48 hours later, cells were infected with SARS-CoV-2. B. qRT-PCR shows SRPK1 mRNA in cells treated with Non-targeting or SRPK1 siRNA transfected A549-ACE2 cells. C. Western Blot of cells that have been treated with Non-targeting or SRPK1 siRNA. D. SARS-CoV-2 viral RNA was measured by qRT-PCR 24h after infection of Non-targeting or SRPK1 siRNA transfected A549-ACE2 cells. E. Cellular viability was measured after A549-ACE2 cells were treated with varying concentrations of SPHINX31 (left axis) and SARS-CoV-2 viral RNA was measured by qRT- PCR 24h after infection of cells that had been treated with varying concentrations of SPHINX31 (right axis). F. Infectious titer was measured by plaque assay of supernatants of cells that were treated with SPHINX31 and infected for 48h. G. Cellular viability was measured after A549-ACE2 cells were treated with varying concentrations of SRPIN340 (left axis) and SARS-CoV-2 viral RNA was measured by qRT- PCR 24h after infection of cells that had been treated with varying concentrations of SRPIN340 (right axis). H. Infectious titer was measured by plaque assay of supernatants of cells that were treated with SRPIN340 and infected for 48h. I. Calu3 cells were infected with varying MOIs and SARS-CoV-2 viral RNA was quantified with qRT-PCR after 24hrs of infection. J. Calu 3 cells treated with DMSO control or SRPIN340 were infected and after 24h post-infection were fixed and stained for DNA with Hoechst and dsRNA using the J2 antibody. Scale bar=150 μm. K. Cellular viability was measured after Calu3 cells were treated with varying concentrations of SRPIN340 (left axis) and SARS-CoV-2 viral RNA was measured by qRT-PCR 24h after infection of cells that had been treated with varying concentrations of SRPIN340 (right axis). L. Infectious titer was measured by plaque assay of supernatants of Calu3 cells that were treated with SRPIN340 and infected for 48h. M. Primary Human Type II Pneumocyte cultures were treated with SRPIN340 for 12h pre-infection, then infected for 24h. Cells were then fixed and stained for SARS-CoV-2, DNA with DAPI, and Surfactant Protein C. Scale bar=100 μm. N. The percent of cells SARS-CoV-2 infected were quantified using 3 Images from 4 different human donors. For all panels *p<0.05 **p<0.001 by an unpaired, student’s t-test.

A549-ACE2 cell lines are an artificial SARS-CoV-2 infection system; we therefore repeated the SRPK1/2 inhibitor experiments with naturally infectible Calu-3 cells (**Figure 3I**) and primary human pneumocytes. Similar to the A549-ACE2 experiments, SARS-CoV-2 infection and replication was significantly inhibited as measured by immunofluorescence for viral replication markers, viral RNA levels, and cell free infectious viral particles (**Figure 3J-N and Figure S4**).

Finally, we were interested in determining if any FDA approved kinase inhibitors could be repurposed to target the phosphorylation of the N protein and thereby inhibit SARS-CoV- 2. Although there are no drugs approved to specifically inhibit SRPK1/2, it is known that the anaplastic lymphoma kinase (ALK) inhibitor Alectinib, which is used clinically to treat non-small-cell lung cancer, also causes significant inhibition of SRPK1/2 (*17*). We therefore treated both A549-ACE2 and Calu-3 cells with Alectinib to investigate inhibition of SARS-CoV-2 infection. Both viral RNA and infectious titer were significantly reduced in a dose dependent manner (**Figure 4A-D**).

**Figure 4.**
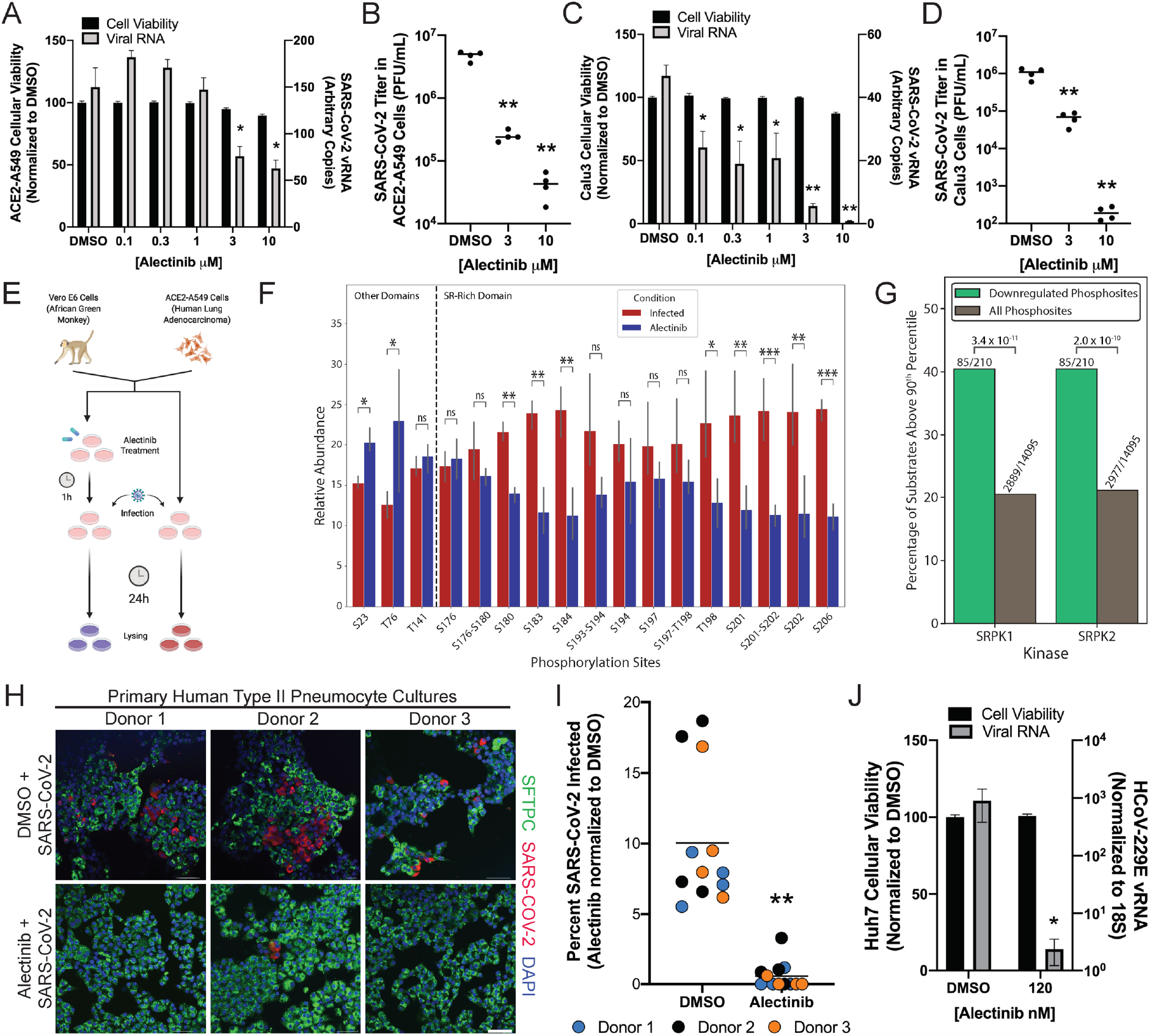
Alectinib can be repurposed to inhibit SARS-CoV-2 infection and reduce N phosphorylation. A. Cell viability was measured after A549-ACE2 cells were treated with varying concentrations of Alectinib (left axis) and SARS-CoV-2 viral RNA was measured by qRT-PCR 24h after infection of cells that had been treated with varying concentrations of Alectinib (right axis). B. Infectious titer was measured by plaque assay of supernatants of cells that were treated with Alectinib and infected for 48h. C. Cell viability was measured after Calu3 cells were treated with varying concentrations of Alectinib (left axis) and SARS-CoV-2 viral RNA was measured by qRT-PCR 24h after infection of cells that had been treated with varying concentrations of Alectinib (right axis). D. Infectious titer was measured by plaque assay of supernatants of Calu3 cells that were treated with Alectinib and infected for 48h. H. Diagram of Alectinib treatment and infection for proteomics and phosphoproteomics analysis. A549-ACE2 and Vero E6 cells were pre-treated with Alectinib (5μM) for 1 h, then infected with SARS-CoV-2 (MOI 0.5) and mock infected for 24h (in biological triplicates each). Proteomics and phosphoproteomics analysis were carried out similarly to the workflow described in Fig. 1A. I. Phosphorylation abundance (normalized by protein level) of the different sites on the N protein, with and without Alectinib treatment in A549-ACE2 cells. The phosphorylation levels of most of the SR-rich domain sites decrease upon Alectinib treatment, while phosphorylation sites outside that domain do not change (or even increase). P-values were computed by moderated t-test and adjusted using Benjamini-Hochberg (BH) correction. n.s. – not significant, *FDR<0.1, **FDR <0.05, ***FDR<0.01. J. The subset of downregulated sites upon Alectinib treatment is enriched for sites that score high for SRPK1 and SRPK2 (above 90^th^ percentile) compared to their abundance among all the detected cellular phosphorylation sites in A549-ACE2 cells. Denoted p-values were computed using Fisher’s exact test. K. Primary Human Type II Pneumocyte cultures were treated with Alectinib for 12h before infection, then infected for 24h. Cells were then fixed and stained for SARS-CoV-2, DNA with DAPI, and Surfactant Protein C. Scale bar=50μm. L. The percent of cells SARS-CoV-2 infected were quantified using 4 Images from 3 different human donors. M. Cell viability was measured after HuH7 cells were treated with Alectinib (left axis) and 229E viral RNA was measured by qRT-PCR 24h after infection of cells that had been treated with Alectinib (right axis). For all panels *p<0.05 **p<0.001 by an unpaired, student’s t-test, unless otherwise stated.

As part of the proteomics and phosphoproteomics experiment described in Figure 1A, biological triplicates of Alectinib pretreated, infected A549-ACE2 and Vero E6 cells were also included together with the control and infected cells (**Figure 4E**). In order to verify that Alectinib was indeed affecting the phosphorylation of the SR-rich domain of the N protein, we analyzed the protein and phosphorylation levels upon treatment, in comparison to infected untreated cells. Viral protein levels were downregulated in the treated cells (**Figure S5A, Table S3**), supporting the hypothesis that Alectinib interferes with the viral life cycle. Examination of the phosphoproteomics data revealed that the vast majority of the SR-rich domain phosphorylation sites were downregulated upon Alectinib treatment, while the phosphorylation levels of sites outside the SR-rich domain did not decrease (**Figure 4F, Figure S5B, Table S1**). Additionally, we scored all the detected (host and viral) phosphorylation sites by SRPK1/2/3 substrate specificity matrices and examined the sites that were downregulated upon Alectinib treatment. The proportion of sites which score high for the SRPK family (scoring above 90^th^ percentile) among the downregulated phosphorylation sites is significantly greater than their proportion among all measured phosphorylation sites (**Figure 4G, Figure S5C**), further confirming that Alectinib inhibits the activity of SRPKs. To demonstrate that Alectinib is capable of inhibiting viral infection outside of immortalized cell lines, we treated our primary human pneumocyte cultures with Alectinib and again observed strong inhibition of the virus (**Figure 4H,I**). These data show that viral replication can be suppressed in some of the most vulnerable populations of lung cells affected during severe COVID-19 disease (*40*), at least *in vitro*, by treatment with the repurposed FDA-approved kinase inhibitor Alectinib. Since the SR-rich domains of N proteins from diverse human coronaviruses are highly conserved (*41, 42*), we next inquired whether the requirement for SRPK1/2 activity might be broadly conserved in this family of viruses. Alectinib treatment of cells infected with the alphacoronavirus 229E (which is only distantly related to betacoronavirus SARS-CoV- 2) inhibited the virus by more than 1,000-fold (**Figure 4J**). These data indicate that the requirement for SRPK1/2 activity is not restricted to SARS-CoV-2 or even betacoronaviruses.

## DISCUSSION

Our study of N protein phosphorylation led to the identification of SRPK1 and SRPK2 as kinases that are critical for the replication of coronaviruses as divergent as 229E and SARS-CoV-2. Further, we provide evidence that the phosphorylation sites in the nucleocapsid protein SR-rich domain and their surrounding sequences are also highly conserved among bat coronaviruses, suggesting that these kinases may also be targetable for pre-pandemic coronaviruses (**Figure S6**). While SRPK1 and SRPK2 are expressed in most human tissues and have been implicated in a number of basal processes including the regulation of transcript splicing, lipid metabolism and cellular stress responses (*43–55*), their activity has also been previously reported as important for the replication of a number of different viruses. These viruses include: hepatitis B virus, human papillomavirus, hepatitis C virus, SARS-CoV, Ebola virus, human cytomegalovirus, and herpes simplex virus-1 (*15, 56–63*). While the mechanisms underlying how SRPK1/2 contribute to the replication of these viruses differ, it is clear that many viruses have evolved to take advantage of these host protein kinases.

Post-translational modification of viral proteins is well understood to be important for the functionality of viral proteins, with phosphorylation chief among them (*7, 64*). At least two recent reports have implicated different kinases including: growth factor receptor (GFR) activated kinases, casein kinase II (CK2), cyclin-dependent kinases (CDKs), and protein kinase C (PKC), as generally important for SARS-CoV-2 replication, but not necessarily directly linked to N protein phosphorylation (*8–10, 34, 65, 66*). Previous work with the related betacoronavirus SARS-CoV has shown that, at least *in vitro*, CDK, GSK, mitogen-activated protein kinase (MAPK), SRPK1, and CK2 can phosphorylate the SARS-CoV N protein (*15, 67*). In this work, while we characterize several of the kinases that are important for phosphorylation of the SARS-CoV-2 N protein SR-rich domain, it is worth noting that other kinases might also play key roles in the phosphorylation of other N protein domains and viral proteins. Understanding the relative contributions of other potential kinases to phosphorylation of different viral proteins and indeed, different domains within the same protein, remains an important area of future study.

Future work will need to establish the functional role of N protein phosphorylation in the SARS-CoV-2 life cycle. A recent study reported that the SARS-CoV-2 N protein phosphorylation affects its protein-protein and protein-RNA interactions through phase separation regulation (*16*). SRPK1 phosphorylation was also reported to affect the ability of the SARS-CoV N protein to multimerize and inhibit host translation, although effects on viral growth were not reported (*15*). Additionally, the growth of both SARS-CoV and mouse hepatitis virus (MHV) have been reported to be suppressed after treatment with GSK-3 inhibitors, presumably at least partially by affecting N protein phosphorylation. It will also be important to define potential kinase redundancy and the effects on viral replication. At least for SARS-CoV-2, it appears that SRPK1/2 are central to the regulation of viral replication. It is worth noting that while our study has focused on the phosphorylation of the SR-rich domain of the viral N protein, we cannot rule out that the viral inhibition phenotype observed may be at least partially due to altered phosphorylation of host proteins or other viral proteins. Additionally, while we have provided data that SRPK1/2 inhibitors are effective in both immortalized and primary human cells, future studies will be required to determine if targeting SRPK1/2 *in vivo* will have a similar magnitude of effect. Although not definitive, favorable outcome of COVID-19 was reported in two cases of patients with non-small cell lung cancer (NSCLC) administered with Alectinib (*68, 69*).

In conclusion, the identification of safe and efficacious anti-COVID-19 therapeutics is of the highest importance due to the ongoing pandemic. Targeting host factors essential for viral replication represents an approach that may help limit the emergence of viral resistance. Our study characterized the activities of several kinases for which at least one FDA-approved inhibitor already exists, can be used to suppress SARS-CoV-2 infection in a variety of *in vitro* systems, and may be repurposed for treating COVID-19. Nevertheless, while Alectinib may have some clinical utility in the short term, continued development of SRPK1/2 specific inhibitors may lead to a class of broadly acting, host-directed antiviral therapeutics that could help combat the current COVID-19, and potentially future, coronavirus pandemics.

## MATERIALS AND METHODS

### Cell culture (Duke)

All cells were obtained from ATCC and grown at 37°C in 5% CO_2_. 293T and A549 and Huh7 cells were grown in DMEM with 10% FBS, Glutamax, and Penicillin/Streptomycin. For ACE2 introduction into A549 cells, both unmodified A549 cells and A549 cells harboring Cas9 were transduced and both of the resulting lines were used for experimentation. Vero E6 cells were grown in MEM media supplemented with Penicillin/Streptomycin, 10% FBS, 1 mM Pyruvate, and 1X MEM NEAA. Calu-3 cells were grown in EMEM with 10% FBS and Penicillin/Streptomycin.

### Cell culture (ISMMS)

Vero E6 cells were obtained from ATCC (CRL-1586). A549 cells stably expressing the SARS-CoV-2 receptor ACE2 were previously described (*71*). All cells were maintained in DMEM supplemented with 10% FBS and Penicillin/Streptomycin 37°C in 5% CO_2_.

### SARS-CoV-2 infections and titering (Duke)

A stock of BEI isolate SARS-CoV-2 USA-WA1/2020 (kind gift of Greg Sempowski) was grown on VeroE6 cells in viral growth media (MEM supplemented with 1% Pen/Strep, 2 % FBS, 1 mM Sodium Pyruvate, and 1x MEM NEAA). Infection was incubated for 1 hr at 37°C. After infection total volume of media was brought to 30 mL. Virus was harvested after 72 hrs of infection. To determine viral titer of stock and after drug treatment, a monolayer of VeroE6 cells was infected with serially diluted virus for 1 hr. Virus was removed and an agar overlay was added to each well (MEM, Penicillin/Streptomycin, 2% FBS, 1 mM Pyruvate, 1x MEM NEAA, 0.3% Sodium Bicarbonate, Glutamax, 0.7% oxoid agar). Plaque assays were incubated for 72 hrs then stained with either 0.1% Crystal Violet in 10% Neutral Buffered Formalin or 0.05% Neutral Red in PBS. All viral infections took place in cell specific media with only 2% FBS.

### HCoV-229E infections and titering

A stock of isolate HCoV-229E VR-740 (ATCC) was grown on Huh7 cells in complete media (DMEM supplemented with 10% FBS, 1% Pen/Strep, Glutamax). Infection was incubated for 1 hr at 37°C. After infection total volume of media was brought to 20 ml. Virus was harvested after 36 hrs of infection. Viral titer of stock was determined via plaque assay on Huh7 cells. A confluent monolayer of cells was infected with serial dilutions of virus in complete media for 1 hr at 37°C. Virus was removed and an agar overlay was added to each well (DMEM, 10% FBS, 1% Pen/Strep, Glutamax, 0.5% oxoid agar). Plaque assays were incubated for 72 hrs then stained with 0.1% Crystal Violet in PBS. Viral infection assays took place in complete media (DMEM, 10% FBS, 1% Pen/Strep, Glutamax).

### SARS-CoV-2 infection for phosphoproteomics (ISMMS)

SARS-CoV-2 isolate USA-WA1/2020 (NR-52281) was deposited by the Center for Disease Control and Prevention and obtained through BEI Resources, NIAID, NIH. SARS-CoV-2 was propagated in Vero E6 cells (ATCC, CRL-1586) in DMEM supplemented with 2% FBS, 4.5 g/L D-glucose, 4 mM L-glutamine, 10 mM Non-Essential Amino Acids, 1 mM Sodium Pyruvate and 10 mM HEPES as previously described (*70*). Virus stocks were filtered and cleared from cytokines and other contaminating host factors by centrifugation through Amicon Ultra-15 100K Centrifugal Filter Units before use. For phosphoproteomic analysis, 6×10^7^ Vero E6 or 6×10^7^ A549-ACE2 cells were treated with 5 µM Alectinib or DMSO in DMEM supplemented with 2% FBS, 4.5 g/L D-glucose, 4 mM L-glutamine, 10 mM Non-Essential Amino Acids, 1 mM Sodium Pyruvate and 10 mM HEPES for 1h at 37°C prior to infection. Cells were subsequently infected with SARS-CoV-2 at an MOI of 0.5 for 24h 37°C. After two washes with PBS and removal of all cell culture media, cell monolayers were lysed in lysis buffer containing 9 M urea, 20 mM HEPES pH 8.0, with 2X phosphatase inhibitors (Cell Signaling Technology #5870).

### Lysis, Digestion, and Preparation for Mass Spectrometry Analysis

Cultured cells were rinsed with phosphate buffer saline (PBS), and scraped into lysis buffer containing 9 M urea, 20 mM HEPES pH 8.0, with 2X phosphatase inhibitors (Cell Signaling Technology #5870). Samples were probe tip sonicated, and subsequently reduced with 5 mM DTT for 50 min at 55 °C, alkylated for 30 min with 10 mM iodoacetamide, and quenched with 5 mM DTT. Samples were diluted to 2M urea with digestion dilution buffer (20 mM pH 8.5 HEPES containing 1mM CaCl_2_) and digested at 37 °C with 20µg of Lysyl Endopeptidase (Wako-Chem) overnight. Samples were then diluted to 1 M urea and digested for 5 hours with 20 µg of trypsin (Pierce). Following digestion, peptides were acidified with trifluoroacetic acid (TFA), centrifuged at 500 x g for 20 min and purified over SepPak C18 columns. Following elution, peptides were quantified with a MicroBCA assay (Thermo Fisher Scientific, San Jose, CA).

### Total Protein Sample Preparation

50µg of peptides from each sample were labeled with isobaric tandem-mass-tag (TMT) 11 plex reagents (Thermo Fisher Scientific, San Jose, CA) in 20 mM pH 8.5 HEPES with 30% acetonitrile (v/v) with 250 ug of TMT reagent. The reaction was quenched for 15 min by adding hydroxylamine to a final concentration of 0.3% (v/v). Samples were combined, dried, purified over SepPak C18 columns, and dried again. Samples were then resuspended in 40 μL of basic reverse phase (bRP) buffer A (10 mM NH_4_HCO_2_, pH10, 5% ACN) and separated on a Zorbax Extended C18 column (2.1 × 150 mm, 3.5 µm, no. 763750-902, Agilent) using a gradient of 10-40% bRP buffer B (10 mM NH_4_HCO_2_, pH10, 90% ACN). 96 fractions were collected and combined into 24 fractions for analysis. Each fraction was dried and desalted over a C18 stop and go extraction tip (STAGE-Tip) prior to analysis by mass spectrometry.

### IMAC Phosphopeptide Sample Preparation

High-Select™ Fe-NTA Phosphopeptide Enrichment Kits from Thermo were used to enrich phosphopeptides from 1mg of peptides for each sample. Following the elution from the IMAC column, enriched samples were dried, labeled with TMT, and bRP fractionated as described above instead with a gradient of 5-40% bRP buffer B. The resulting 24 fractions were desalted over a C18 STAGE-Tip.

### LC–MS Analysis of Total Protein Fractions

Samples were analyzed on an Orbitrap Fusion Lumos mass spectrometer (Thermo Fisher Scientific, San Jose, CA) coupled with a Proxeon EASY-nLC 1200 liquid chromatography (LC) pump (Thermo Fisher Scientific, San Jose, CA). Peptides were separated on a 100 μm inner diameter microcapillary column packed with ∼40 cm of Accucore150 resin (2.6 μm, 150 Å, ThermoFisher Scientific, San Jose, CA). For each analysis, we loaded approximately 1 μg onto the column. Peptides were separated using a 2.5h gradient of 6–30% acetonitrile in 0.125% formic acid with a flow rate of 550 nL/min. Each analysis used an SPS-MS3-based TMT method (*72, 73*), which has been shown to reduce ion interference compared to MS2 quantification (*74*). The scan sequence began with an MS1 spectrum (Orbitrap analysis, resolution 60,000; 350-1400 m/z, automatic gain control (AGC) target 4.0 × 10^5^, maximum injection time 50 ms). Precursors for MS2/MS3 analysis were selected using a Top10 method. MS2 analysis consisted of collision-induced dissociation (quadrupole ion trap; AGC 2.0 × 10^4^; normalized collision energy (NCE) 35; maximum injection time 120 ms). Following acquisition of each MS2 spectrum, we collected an MS3 spectrum using a method in which multiple MS2 fragment ions are captured in the MS3 precursor population using isolation waveforms with multiple frequency notches (*73*). MS3 precursors were fragmented by HCD and analyzed using the Orbitrap (NCE 65, AGC 3.5 × 10^5^, maximum injection time 150 ms, isolation window 1.2 Th, resolution was 50,000 at 200 Th).

### LC–MS Analysis of Phosphopeptide Enriched Fractions

For phosphopeptide analysis the same MS and HPLC instruments were used as described above. For each analysis approximately 500 ngs of enriched peptides were loaded on the column and run over a 120-minute gradient of 2-32% acetonitrile in 0.125% formic acid with a flow rate of 400 nL/min. MS1 spectra were collected in the Orbitrap at a resolution of 60,000 with a scan range of 300-1500 m/z using a AGC target of 4.0e5 with a maximum inject time of 25ms. Peptides for MS2 analysis were isolated using the quadrupole with an isolation window of 0.5 m/z. MS2 spectra were generated using Higher-energy collision dissociation (HCD) with a collision energy of 40%. Fragments were collected in the Orbitrap at a resolution of 50,000 with a first mass of 110m/z, an AGC target of 5.0e4 and a maximum injection time of 200ms.

### Total Protein Data Processing and Analysis

Mass Spectra were processed using a Comet-base software pipeline (*75, 76*). Resulting data were searched with a fully tryptic database containing both human Swissprot consensus entries plus isoforms downloaded Feb 2020, and SARS-CoV-2 pre-release entries from Uniprot (June 2020), allowing for a static modification of lysine and N-termini with TMT (229.1629 Da) and carbamidomethylation (57.0215 Da) of cysteine, along with variable oxidation (15.9949 Da) of methionine. Searches were performed using a 20 ppm precursor ion tolerance, the product ion tolerance was set to 1.0 Th. Peptide-spectrum matches (PSMs) were adjusted to a 1% false discovery rate (FDR) using previously described linear discriminant analysis (*77, 78*). Filtered PSMs were collapsed to a final protein-level FDR of < 1%. Protein assembly was guided by principles of parsimony to produce the smallest set of proteins necessary to account for all observed peptides (*75*). For TMT-based reporter ion quantitation, we extracted the summed signal-to-noise (S/N) ratio for each TMT channel and found the closest matching centroid to the expected mass of the TMT reporter ion. MS3 spectra with TMT reporter ion summed signal-to-noise ratios less than 100 were excluded from quantitation (*79*).

### Phosphorylation Data Processing and Analysis

When searching phosphorylation data, a variable modification for phosphorylation (79.9663 Da) was allowed on serine, threonine, and tyrosine. Searches were performed using a 20 ppm precursor ion tolerance, the product ion tolerance was set to 0.02 Th. Linear discriminant analysis (LDA) was performed to set a PSM FDR of < 1 %. Filtered PSMs were collapsed to a final protein-level FDR of < 1%. Phosphorylation sites were evaluated using an AScore method. Sites with an AScore > 13 were considered localized (*80*). PSMs with TMT reporter ion summed signal-to-noise ratios less than 100 were excluded from quantitation.

### Quantification and normalization of TMT data

For each entry (protein or phosphorylation site), the level in each TMT channel was calculated as the log-transformation of the ratio to the median value of all 9 TMT channels (3 mock, 3 infected and 3 treated) of that entry. Then, every sample was median-normalized so the log-TMT-ratio is centered at zero. Phosphorylation levels were normalized by protein levels by subtracting the log-TMT-ratio of the corresponding protein from the log-TMT-ratio of the phosphorylation site.

### Differential expression analysis

Differential expression analysis of proteins and phosphorylation sites was done using Limma v3.42 package in R (*81*). For protein levels, log-TMT-ratio was used as input data, and for phosphorylation sites – the protein-normalized log-TMT-ratio was used. Since the log-TMT-ratio is normally distributed, no voom-normalization was applied. Unequal variance between samples was taken into account by using the “arrayWeight” function before fitting the model. P-values were computed using moderated t-test, and adjusted p-values (FDR) were calculated using the Benjamini–Hochberg (BH) correction. Significance for differential expression was determined as adjusted P-value<0.1.

### Kinase substrate specificity assays

Reagents used for the peptide library experiments include: Kinase substrate library (Anaspec). Streptavidin conjugated membranes (Promega). All the recombinant kinases (SRPK1/2/3, GSK-3α/β and CK1A/E were obtained from SignalChem.

To determine the substrate motifs, we performed *in vitro* phosphorylation assays with recombinant kinases on an oriented peptide array library of design:

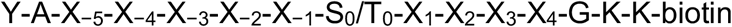

in the presence of ATP[γ-^32^P]. Reactions were carried out in their designated buffers plus 20 μm ATP and 0.4 μCi of (33 nm) [γ-^32^P]ATP) at 30°C for 90 min. The peptides were spotted onto streptavidin-coated filter sheets (Promega SAM^2^ biotin capture membrane) and visualized by phosphorimaging on Typhoon FLA 7000. Detailed information on the protocol is provided elsewhere (*30, 31*).

### Matrix processing and substrate scoring

The matrices were normalized by the sum of the 17 randomized amino acids (all amino acids expect for serine, threonine and cysteine), to yield a position specific scoring matrix. The serine, threonine and cysteine columns were scaled by their median to be 1/17. For scoring substrates, the values of the corresponding amino acids in the corresponding positions were multiplied and scaled by the probability of a random peptide:

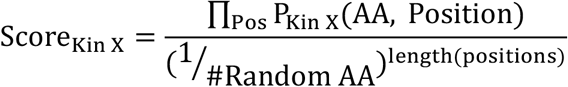

For the percentile score of a substrate by a given kinase, we first computed the *a-priori* score distribution of that kinase by scoring all the reported S/T phosphorylation sites on PhosphoSitePlus (*82*) (downloaded on January 2020) by the method discussed above. The percentile score of a kinase-substrate pair is defined as the percentile ranking of the substrate within the score distribution of the given. This value is being used when analyzing all the detected phosphorylation sites (viral and host) for kinase enrichment.

### Evolutionary conservation analysis

All reference proteomes of the Coronaviridae family of viruses available on Uniprot were downloaded (82 in total, see **Supplementary Table S3**). A BLAST search was performed over each proteome using the SARS-CoV-2 nucleocapsid protein as the query; the top hit of each virus was taken to be the nucleocapsid. Each designated nucleocapsid was then individually aligned to the SARS-CoV-2 nucleocapsid using MUSCLE. Using these alignments, the percentage identity was then calculated for each position along the SARS-Cov-2 nucleocapsid. All statistical analyses between different conservation of positions were performed using the Mann-Whitney U test.

### Kinase enrichment analysis

The phosphorylation sites detected in this study were scored by SRPK1/2/3 substrate specificity matrices and their ranks in the known phosphoproteome score distribution was determined as described above (percentile score). For every assessed kinase, phosphorylation sites that ranked within the top 10 percentile within the score distribution were counted as biochemically favored sites for that kinase. Towards assessing SRPK kinase motif enrichment, we compared the percentage of biochemically favored sites in the downregulated peptides (log_2_Fold Change of −0.75 and below with an FDR of >= 0.1) to the percentage biochemically favored sites within the set of all detected sites in this study. Statistical significance was determined using Fisher’s exact test.

### Sequence logos

Normalized substrate specificity matrices were scaled to represent relative probability (each position sum up to 1), and probability sequence logos were generated using ggseqlogo v0.1 (*83*) package in R.

### Plasmids

The ACE2-pLEX overexpression plasmid was generated by cloning a synthesized gBlock of the ACE2 ORF (Ref. seq. NM_001371415.1) into the pLEX plasmid via HiFi DNA Assembly (NEB Cat# M5520AA) and subsequent bacterial transformation. Bacterial colonies were then selected and plasmid DNA isolated using the Genejet Plasmid Miniprep kit (Thermo Fisher Cat# K0503). Plasmids were then sequenced via Sanger sequencing to confirm successful cloning of the ACE2 ORF. Lentiviruses were packaged as per standard protocols with a VSV-G envelope protein, A549 or A549-Cas9 cells were transduced and selected with puromycin at 2μg/mL.

### siRNA treatment of cells

A549-ACE2 cells were treated with 30μM siRNA (SRPK1: Horizon M-003982-02-0010; Non-targeting: Thermo 4390843) following the HiPerfect (Qiagen) Fast-Forward protocol and plated in 6 well plates. After 2 days, cells were trypsinized and replated in 24 well plates and re-transfected with siRNA (50μM) using the same protocol. After 2 additional days, cells were infected at MOI 0.005 for 1h in 250μl DMEM+2% FBS. After 1h of infection, 250μl DMEM+2% FBS was added to the inoculum and cells were incubated for 24h. 24h post-infection the media was removed and cells were lysed with 500μl TriZol. RNA was purified according to protocol below.

### Quantitative Reverse Transcription PCR

Cells were resuspended in TRIzol Reagent (Thermo Fisher Cat #15596018) and total RNA was isolated by either phase separation with chloroform and isopropanol or using the Zymo Direct-zol RNA Miniprep kit (Zymo Cat #R2050). One-step qRT-PCR was performed using the Invitrogen EXPRESS One-Step Superscript qRT-PCR kit (Thermo Fisher Cat #11781200) and the commercial Taqman probe ACE2 (Hs01085333_m1). The SARS-CoV2 primer/probe set were synthesized using IDT DNA based on the sequences provided by the CDC “Research Use Only 2019-Novel Coronavirus (2019-nCov) Real-time RT-PCR Primers and Probes” set N1. For qRT-PCR of SRPK1, commercial Taqman probe (Hs00177298_m1) was used. For RNA extractions of 229E infected cells, RNA was extracted using NEB Monarch total RNA miniprep kit (T2010S). To quantify viral RNA we used commercial taqman probe CoV_229E (Vi06439671_s1). Reactions were cycled on the Applied Biosystems QuantStudio 3 Real-Time PCR System and analyzed using QuantStudio software version v1.4.1. RNA was normalized to the endogenous 18S primer-probe set (Thermo Fisher Cat #4319413E).

### Western blots

Cells were trypsinized, removed from plates, and pelleted via centrifugation (3,000*g* for 4 minutes). Cell pellets were washed once with PBS. Cells were lysed via resuspension in RIPA buffer (10 mM Tris-HCl pH 7.5, 1 mM EDTA pH 8.0, 1% Triton X-100, 0.1% sodium deoxycholate, 140 mM NaCl, 0.1% SDS) and incubation at 4 °C for 15 minutes. Chromosomal DNA was sheared by passing lysates through an insulin syringe and cellular debris was removed via centrifugation (21,100*g* for 10 minutes at 4 °C). Total protein concentration of cellular lysates was determined via Bradford assay and all samples were normalized to 1 µg/µL. Cellular lysates were separated by SDS-PAGE using 15-20 µg total cellular protein per lane (4-20% Mini-PROTEAN TGX gels (BioRad), 120 V for 1 hour). Proteins were subsequently transferred to nitrocellulose membranes (90 V for 1 hour at 4 °C). Nitrocellulose membranes were blocked for at least 3 hours using PBST containing 5% milk. For detection of SRPK1, membranes were incubated with primary antibody (BD cat. no. 611072) at 0.025 µg/mL in PBST containing 5% milk overnight at 4 °C. For detection of GAPDH, membranes were incubated with primary antibody (Cell Signaling, cat. no. 2118S) at a dilution of 1:10,000 in PBST containing 5% milk overnight at 4 °C. Anti-mouse (Invitrogen cat. no. A16072) and anti-rabbit (Thermo Scientific cat. no. A16104) secondary antibodies were used at 1:20,000 and 1:10,000 dilutions, respectively, in PBST containing 5% milk for 1 hour at room temperature. Blots were developed using Clarity Max ECL substrate (BioRad).

### Crystal violet staining

To visualize CPE of A549 cells expressing ACE2 or not, cells were infected with a MOI of 0.1 for 1h at 37°C. After 1h incubation 0.5 ml DMEM with 2% FBS was added to each well and infection was allowed to continue for 72h. After infection, the monolayer was stained with 0.1% Crystal violet in 10% Neutral Buffered Formalin for 15 minutes. Stain was then removed and cells were rinsed with ultra-pure water and then imaged.

### SRPK inhibitor treatment assays

Three different inhibitors of SRPK1 and SRPK2 were used to assay effects on viral replication, SRPIN340, SPHINX31, and Alectinib (MedChem Express). SRPIN340 and SPHINX were suspended in DMSO at 1000X the final concentration indicated, Alectinib at 500X. They were then diluted to the indicated concentrations in cell specific media. Cells were treated with drug 12 hours before infection. Cells were then infected for 1 hour as described above. After infection, media with 2% FBS + drug or DMSO control was added to each well (qRT-PCR and Calu-3 microscopy) or inoculum was removed and media with 2% FBS + drug or DMSO control was added to each well (infectious titer quantification).

### Cytotoxicity Assays

A549-Cas9 ACE2, Calu-3, or Huh7 cells were treated with the indicated concentrations of SRPIN-340, SPHINX-31, or Alectinib in a consistent volume of vehicle (DMSO). 48 hours after treatment, samples were processed according to the CellTiter-Glo (Promega) protocol and luminescence was determined using a luminometer.

### Calu-3 Infection and immunofluorescence assays

For infections of Calu-3 cells for immunofluorescence assays, cells were treated with DMSO or 53 μM SRPIN340 12h before infection. To infect Calu-3 cells media was removed and cells were washed with 0.5 mL PBS. Virus was added at MOI 2.5 and cells were infected at 37°C for 1h. After infection, 200 μL media + 2x SRPIN340 or DMSO was added to cells and incubated at 37°C for 24h. To fix cells, the culture plate was submerged in 10% neutral buffered formalin for 2h. NBF was removed and cells were placed in PBS. For immunostaining cells were permeabilized with 0.1% Triton X-100 and blocked in PBS+ 5% BSA + 0.1% Tween-20. Cells were stained with either an anti-dsRNA antibody (J2, Sigma MABE1134) at a 1:125 dilution, or an anti-SARS-CoV-2 Spike RBD protein (ProSci #9087) at a 1:150 dilution. Alexa fluor 488 or 594 conjugated secondary antibodies were used at a 1:1000 dilution. DNA was stained with Hoechst 33342 dye at a 1:10,000 dilution in PBS. Images were taken on Bio-Rad Zoe fluorescent cell imager.

### In vitro SRPK1/2 phosphorylation assays

2 μM recombinant N protein was incubated with recombinant kinases SRPK1 (80 nM), GSK-3A (5 nM), and/or CK1ε (30 nM) in Kinase Buffer 1 (SignalChem) for 15 minutes at 30 degrees C. Reactions were terminated with the addition of SDS loading buffer. The reactions were run phos-tag gel and blotted with monoclonal anti-N protein antibody (GeneTex). For autoradiography, the reactions were supplemented with γ[^32^P]-ATP and were run on an SDS-PAGE. The N protein bands (48 kDa) were excised and radioactivity was measured by Typhoon FLA 7000.

### Human lung dissociation, primary type two pneumocytes purification, culturing, and immunostaining

Human lung tissues were washed with PBS containing 1% Antibiotic-Antimycotic and cut into small pieces. Then samples were digested with enzyme mixture (Collagenase type I: 1.68 mg/ml, Dispase: 5U/ml, DNase: 10U/ml) at 37°C for 1 h with rotation. The cells were filtered through a 100 µm strainer, rinsed with 15 ml DMEM/F12+10% FBS medium and centrifuged at 450x g for 10 min. The supernatant was removed and the cell pellet was resuspended in red blood cell lysis buffer for 10min, washed with DMEM/F12 containing 10% FBS and filtered through a 40 µm strainer. Total cells were centrifuged at 450xg for 5 min at 4°C and the cell pellet was processed for AT2 purification.

For human type two pneumocytes purification, approximately 2-10 million total human lung cells were resuspended in MACS buffer and blocked with Human TruStain FcX (Biolegend cat# 422032) for 15min at 4°C followed by HTII-280 antibody (Terrace Biotech, TB-27AHT2-280) staining for 1 h at 4°C. The cells were washed twice with MACS buffer followed by incubation with anti-mouse IgM microbeads for 15 min at 4°C. The cells were loaded into the LS column and collected magnetically. AT2 cells (3×10^3^) were resuspended in serum free medium, mixed with an equal amount of Matrigel (Corning Cat# 354230) and plated as drops on 6 well plates. The medium was changed every three days. Alveolospheres were passaged every 14 days. For virus infection experiments human AT2 cells were seeded at 10 x 10^4^ cells per insert on 5% matrigel coated Transwell with 0.4-μm pore-sized inserts (Corning). Cells were treated with DMSO or 53 μM SRPIN340 12h before infection with media on the top and bottom of transwell. To infect cells, media was removed from top of transwell and cells were washed with 200 μl PBS. Cells were then infected at a MOI of 1 for 1h. Virus was then removed, and cells were cultured for 24h. To fix cells and remove from BSL3, plates and wells were submerged in 10% Neutral buffered formalin for 2h. NBF was removed and cells were placed in PBS until staining.

For immunostaining of the cultures, the membrane from the transwell insert was cut out with a scalpel and washed with PBS, permeabilized in PBST (0.1% Triton X-100 in PBS), and incubated with blocking buffer (1% BSA in PBST) for 1 h at room temperature. Samples were incubated with primary antibodies: Prosurfactant protein C (1:500, Milipore, cat# AB3786) and SARS-CoV-2 (1:500, Genetex cat# GTX632604) in blocking buffer at 4°C overnight. Membranes were then washed 3 times in PBST, incubated with secondary antibodies in blocking buffer for 1 h at room temperature followed be three washes with PBST and mounted using Fluor G reagent with DAPI. All confocal images were collected using Olympus Confocal Microscope FV3000 using a 40X objectives.

## ACKNOWLEDGMENTS

The authors would like to acknowledge experimental support from, and helpful discussion with, Hal Bogerd and Bryan Cullen. This research was supported by funding from the Pershing Square Foundation (JB and LCC) and RO1 GM51405-28 (J.B.) NSH is partially funded by the Defense Advanced Research Projects Agency’s (DARPA) PReemptive Expression of Protective Alleles and Response Elements (PREPARE) program (Cooperative agreement #HR00111920008). The views, opinions and/or findings expressed are those of the authors and should not be interpreted as representing the official views or policies of the U.S. Government. JDT was partially supported by T32- CA009111. The following reagent was deposited by the Centers for Disease Control and Prevention and obtained through BEI Resources, NIAID, NIH: SARS-Related Coronavirus 2, Isolate USA-WA1/2020, NR-52281. Biocontainment work was performed in the Duke Regional Biocontainment Laboratory, which received partial support for construction from the National Institutes of Health, National Institute of Allergy and Infectious Diseases (UC6-AI058607). Biocontainment work was also performed at Icahn School of Medicine at Mount Sinai.

