## Supplemental Data for "The FDA-approved drug Alectinib compromises SARS-CoV-2 nucleocapsid phosphorylation and inhibits viral infection in vitro"

### Supplemental Figures and Tables

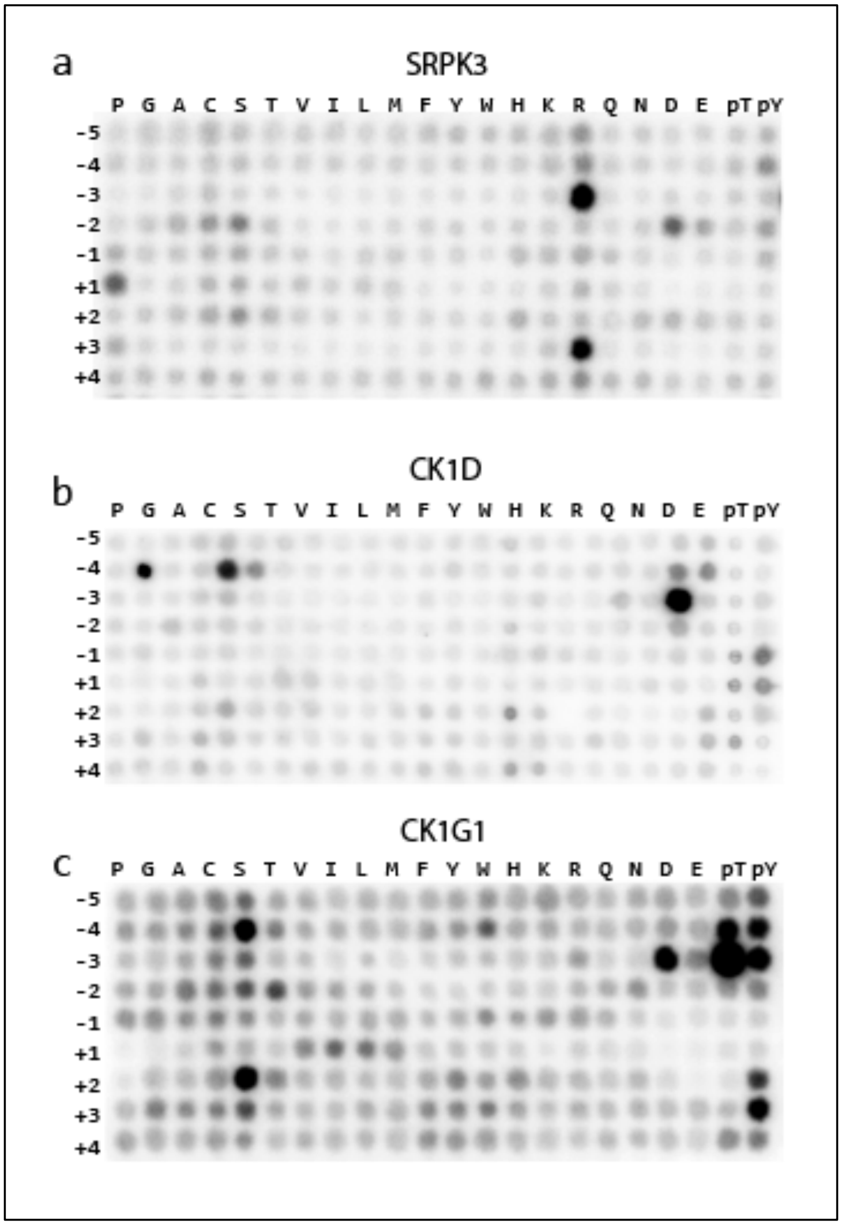


**Supplemental Figure 1. Biochemical substrate specificities of SRPK3, CK1D and CK1G1.** (a) Similar to SRPK1/2, also SRPK3 is selective for arginines at the -3 and +3 positions, serines at the -2 and +2 positions, and proline at the +1 position. (b-c) Similar to CK1A/ε, also CK1D and CK1G1 are selective for phosphoserine and phosphothreonine at position -3 and for serine at position -4.

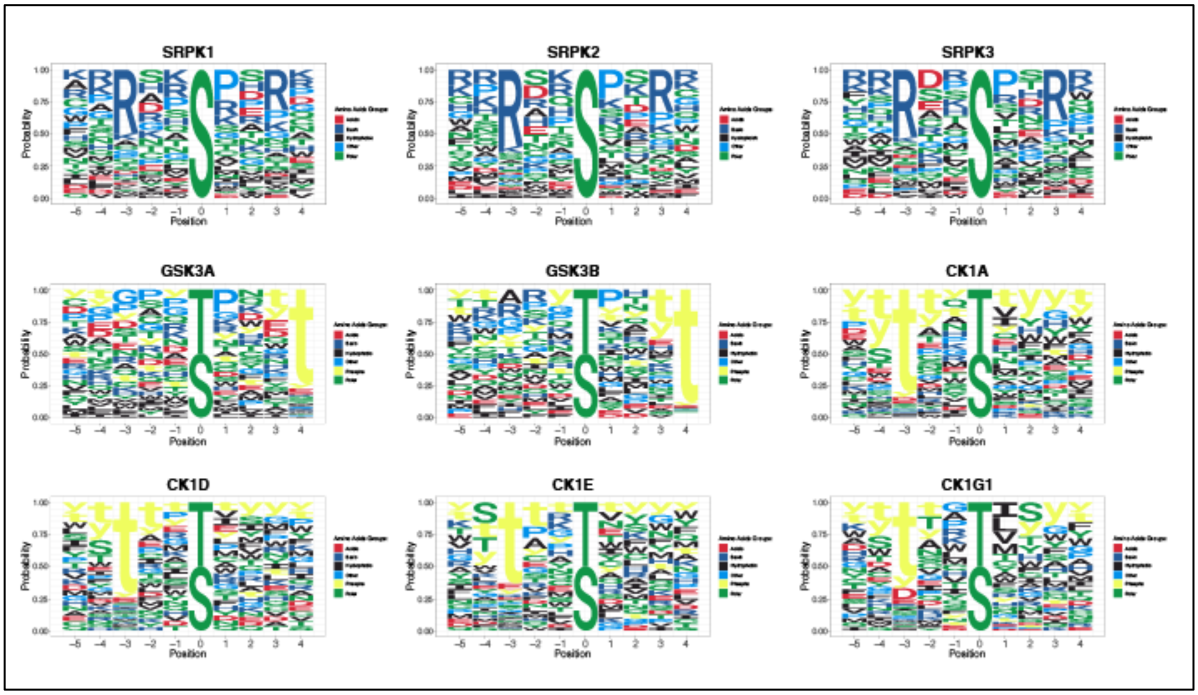


**Supplemental Figure 2. Sequence Logos of SRPK, GSK3 and CK1 families, based on their biochemical substrate specificity matrices.** The values of the normalized substrate specificity matrices were converted into relative probability and plotted as a sequence logo. For GSK3 and CK1 families, phosphorylated residues were also included.

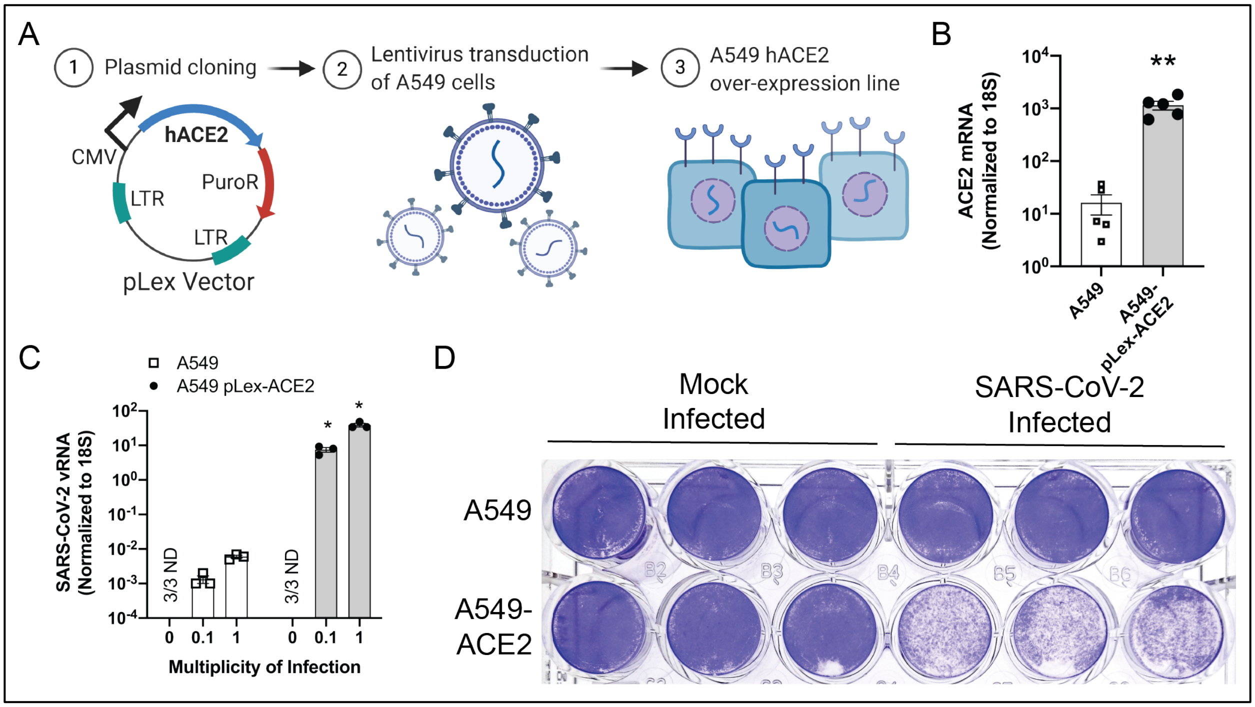


**Supplemental Figure 3. Development and validation of an A549-ACE2 line for SARS-CoV-2 infection.** (a) Scheme for construction of the A549-ACE2 cell line. (b) Quantification of *Ace2* mRNA by qRT-PCR in the parental A549 cells or A549 cells transduced with pLex-ACE2 lentivirus, n=5 for both treatment groups. (c) Quantification of SARS-CoV-2 RNA after 24 hrs infection of A549 or A549-ACE2 cells (n=3) at different MOIs. (d) Crystal violet staining of A549 or A549-ACE2 cells after 72 hrs of SARS-CoV-2 infection with MOI=0.1.

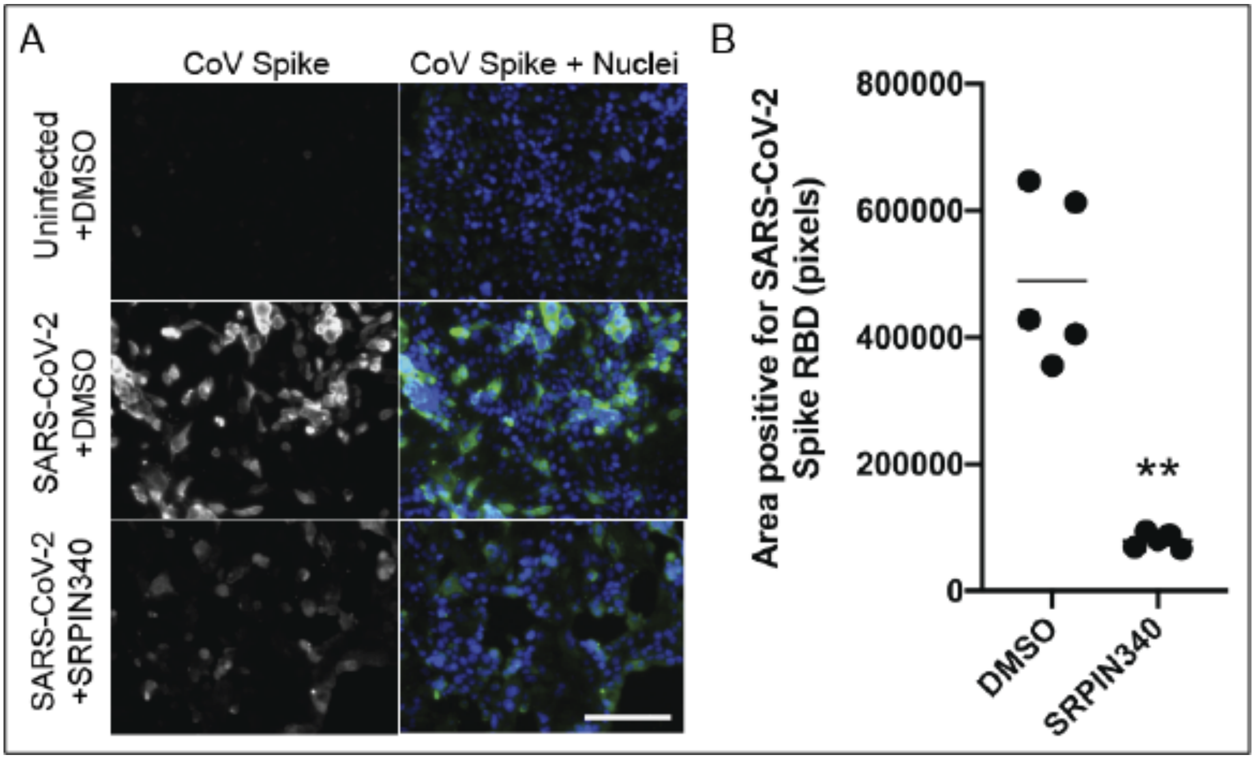


**Supplemental Figure 4. Microscopy for SARS-CoV-2 Spike in Calu-3 cells with and without SRPIN340 treatment.** (a) Calu-3 cells were uninfected or infected with SARS-CoV-2, MOI=3 after treatment with DMSO control or 53 μM SRPIN340. Nuclei were stained with Hoechst (blue) and SARS-CoV-2 Spike protein is in green. Scale bar represents 150 μm. (b) SARS-CoV-2 Spike protein staining was quantified via ImageJ from the experiment described in (a), n= 5 images per treatment.

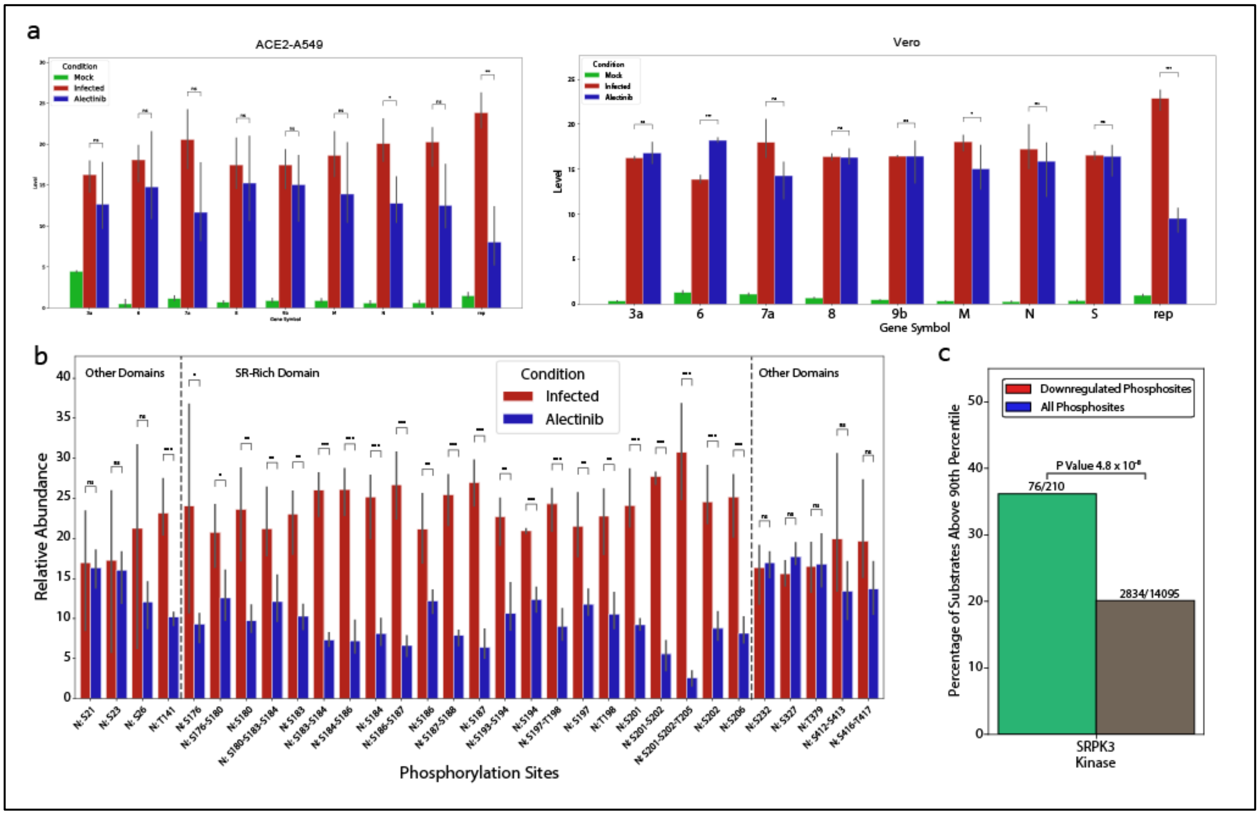


**Supplemental Figure 5. Proteomics and phosphoproteomics analysis of Alectinib pre-treated, infected ACE2-A549 and Vero E6 cells.** (a) Viral protein levels in ACE2-A549 (left) and Vero (right). Upon Alectinib treatment, the replicase polyprotein (ORF1ab) is significantly downregulated in both cell lines. P-values were computed by moderated t-test and adjusted using Benjamini-Hochberg (BH) correction. n.s. – not significant, *FDR<0.1, **FDR <0.05, ***FDR<0.01. (b) Phosphorylation sites levels (normalized by protein level) in Vero cells upon Alectinib treatment. The phosphorylation levels of most of the SR-rich domain sites decrease upon Alectinib treatment, while most of the phosphorylation sites outside that domain do not change. P-values were computed by moderated t-test and adjusted using Benjamini-Hochberg (BH) correction. n.s. – not significant, *FDR<0.1, **FDR <0.05, ***FDR<0.01. (c) SRPK3 enrichment analysis for the downregulated phosphorylation sites upon Alectinib treatment. Denoted p-values were computed using Fisher’s exact test.

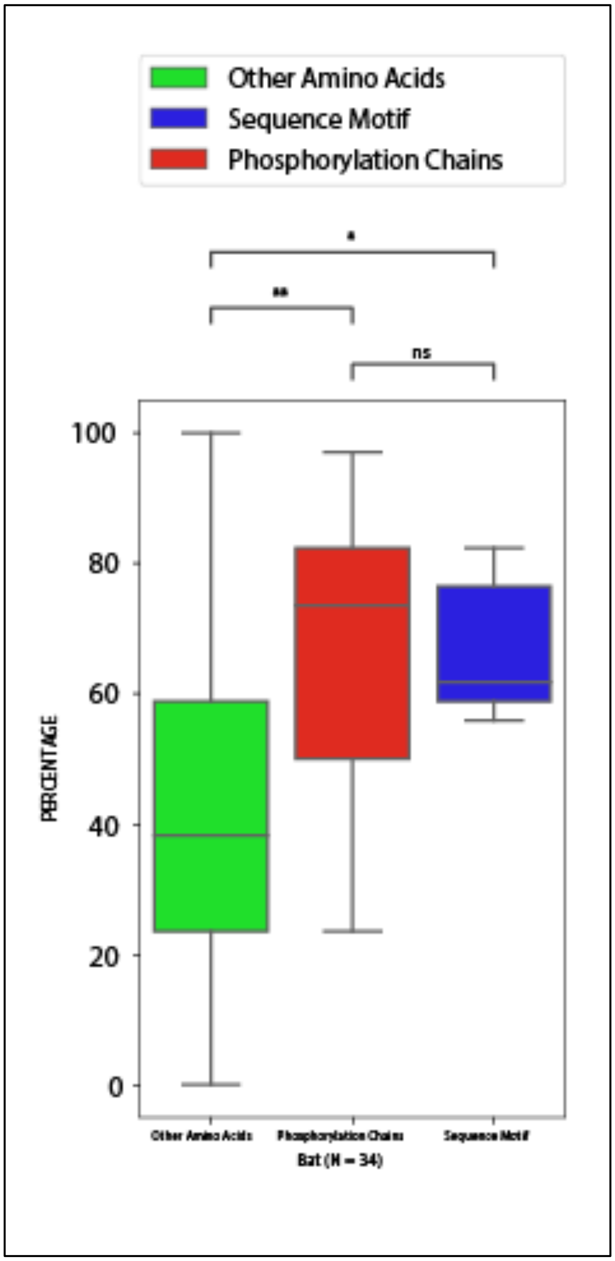


**Supplemental Figure 6. Evolutionary conservation across 34 different bats-specific coronaviruses.** Also in bats-specific coronaviruses, amino acids predicted to be essential for substrate specificity of the priming sites are as conserved as the phosphorylation sites themselves, and are more conserved than the other amino acids in the SR-rich domain, suggesting that these kinases may also be targetable for pre-pandemic coronaviruses. Mann-Whitney U test: n.s. – not significant, *p<0.05, **p<0.01.
